# The chromosome level genome of the Blueberry Stem Gall Wasp, *Hemadas nubilipennis* (Hymenoptera: Ormyridae) on lowbush blueberry (*Vaccinium angustifolium*) reveals repeat-driven expansion

**DOI:** 10.64898/2026.09.21.753281

**Authors:** Y. Miles Zhang, Adam J. Kranz, Renee L. Corpuz, Soham Mukhopadhyay, Ellen O. Martinson, Michael Sergeant, Glen Ray Hood, Scott M. Geib, Sheina B. Sim

## Abstract

Gall-inducing insects are important models for studying plant-feeding lifestyle evolution, but genomic resources remain scarce. We present the first chromosome-level genome for the blueberry stem gall wasp (*Hemadas nubilipennis*), a native North American species now reaching outbreak densities on cultivated highbush blueberries. The 1.08 Gb genome (N50=218 Mb) is the second largest in Chalcidoidea, with size variation driven by transposable element proliferation (R²=0.96). Gene-body methylation is conserved and correlates with gene density. Mitochondrial COI sequences reveal 4.3–5.4% divergence among geographically separated populations, suggesting a cryptic species complex. We also assemble a near-complete *Wolbachia* genome (Supergroup A) encoding parthenogenesis-linked effectors. These resources establish foundations for population genomics, taxonomic revision, and applied management, while providing insights into genome architecture, epigenetics, and symbiont interactions, offering essential tools for distinguishing cryptic species and understanding this emergent pest’s biology.

## Introduction

The family Ormyridae (Hymenoptera: Chalcidoidea) comprises six genera and 153 species (UCD Community 2023; Hanson et al. 2025), associated with insect-induced galls as inducers or parasitoids (Hanson et al. 2025). Recent phylogenomic work reclassified Ormyridae to include gall-inducing taxa (van Noort et al. 2024). The blueberry stem gall wasp (BSGW), *Hemadas nubilipennis* (Figure 1A), was moved from Pteromalidae and is now the sole member of Hemadinae (van Noort et al. 2024; Hanson et al. 2025).

**Figure 1.**
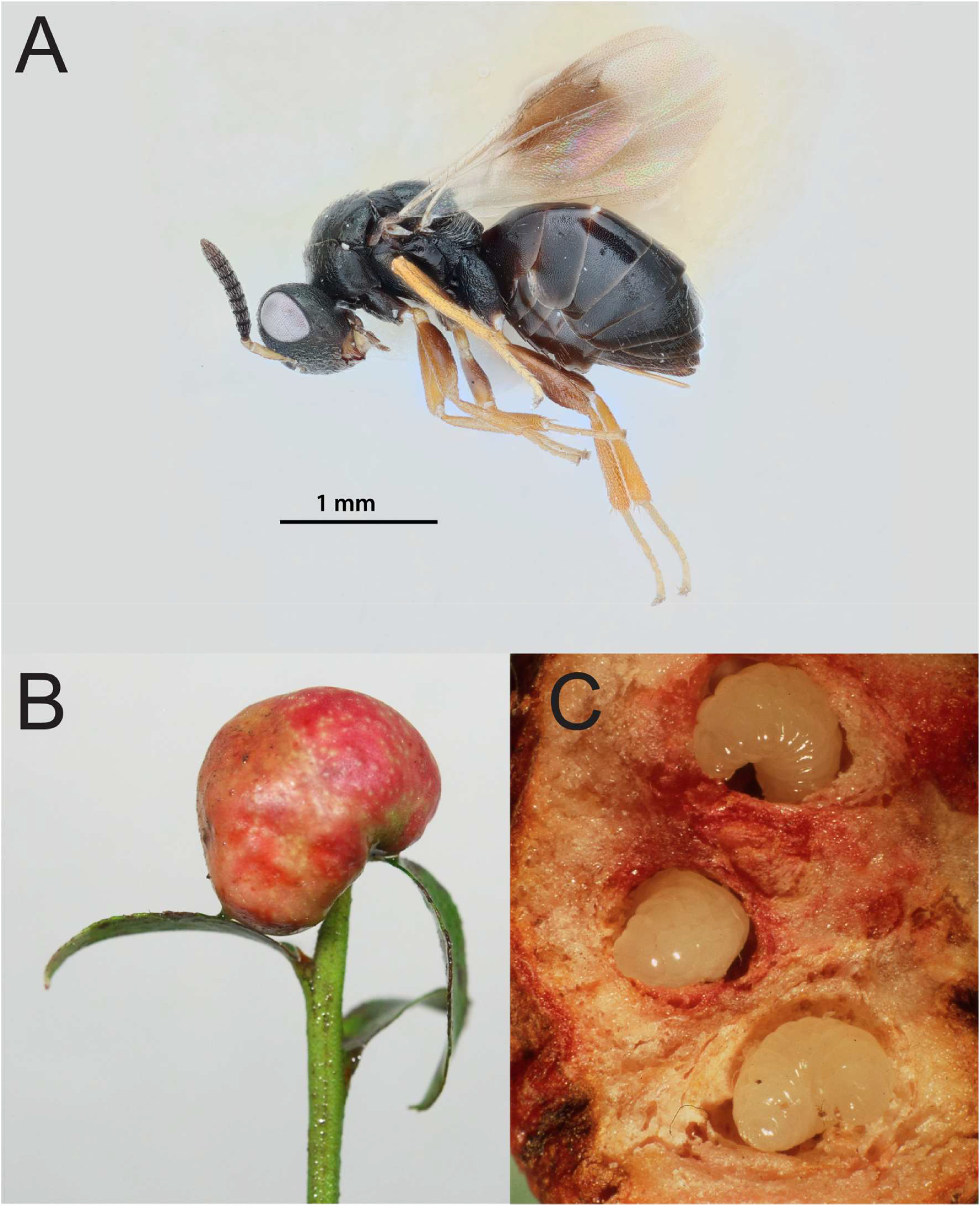
Blueberry stem gall wasp (*Hemadas nubilipennis*) on lowbush blueberry (*V. angustifolium*). **A)** Lateral habitus of adult female. **B)** Gall, photo by Joseph Shorthouse. **C)** Mature larvae, photo by Joseph Shorthouse.

BSGW is native to the midwestern and eastern U.S. and eastern Canada, inducing galls on native highbush (*Vaccinium corymbosum*), lowbush (*V. angustifolium*), and cultivated highbush blueberry (Isaacs et al. 2020; Teresi et al. 2025) (Figure 1B, 1C). Recently, BSGW reached outbreak densities in southwestern Michigan (Isaacs et al. 2020), where blueberry production is valued at $132 million annually (Wise and Garcia-Salazar 2023). Damage occurs after oviposition; females stab the meristem, redirecting nutrients to gall tissue and terminating shoot growth, reducing fruit production (Shorthouse et al. 1986; West and Shorthouse 1989; Hayman et al. 2003). Given the pest’s broad range and the U.S. blueberry industry generating >$4.5 billion annually, genomic resources are urgently needed.

The Michigan outbreak is unusual because BSGW is not invasive (Bertelsmeier et al. 2025). Historically, populations remained at low densities across its range, including in Michigan. Researchers speculate that natural enemy suppression, insecticide shifts, or host resistance variation may contribute (Isaacs et al. 2020; Fanning and Isaacs 2020; DeVisser et al. 2023; Teresi et al. 2025), but the drivers remain unknown. Whether BSGW populations on different hosts represent cryptic species is unknown, but three lines of evidence suggest this. First, highbush and lowbush blueberries occupy distinct niches with largely allopatric ranges and highly divergent growth forms. Second, host-associated fitness traits differ: galls on lowbush are smaller, and populations are strongly female-biased (98–100%), whereas highbush and cultivated populations are more equitable (60–80% female; Shorthouse et al. 1990; G.R. Hood and M. Sergeant, unpublished). Third, host-associated populations harbor different natural enemy communities (Shorthouse et al. 1990; Isaacs et al. 2020). These ecological differences suggest divergent selection that could promote cryptic speciation (Forbes et al. 2017; Martinson et al. 2026).

Here, we present the first chromosome-level genome for *H. nubilipennis* from a lowbush-associated population. We chose this lineage because: (i) it is the best-characterized natural history system (Shorthouse et al. 1986, 1990; Wise and Shorthouse 1990; Hayman et al. 2023); (ii) its extreme female bias enables investigation of reproductive endosymbionts like *Wolbachia* (Fricke and Lindsey 2024); (iii) it provides a baseline for comparing host-associated divergence and cryptic speciation; and (iv) it establishes an evolutionary baseline for identifying genomic changes associated with adaptation to cultivated systems. Using this reference, we test whether its exceptionally large size is driven by transposable element proliferation, a pattern linked to phytophagy, and characterize its methylation landscape and symbiont content to understand its reproductive mode and cryptic diversification.

## Materials and methods

### Source material

In late winter 2025, about two months prior to natural *H. nubilipennis* adult emergence in the field, we collected galls containing overwintering larvae from lowbush blueberry at a single site in Lake County in northwestern Michigan near Wolf Lake (44.02427 N, −85.82244 W) used for genomic sequencing. The galls were placed in a cooler, transported to the laboratory at Wayne State University, and stored in a cooler at 4°C. In late May, galls were removed from the cooler and placed in clear, 470-mL plastic cups that were monitored twice daily for emergence. Upon emergence, individuals were taxonomically identified, sexed, and flash frozen in liquid nitrogen and then stored at −80°C. A total of 10 female *H. nubilipennis* were shipped, overnight, on dry ice to the Daniel K. Inouye U.S. Pacific Basin Agricultural Research Center in Hilo, Hawaiʻi. Lateral habitus image of the adult wasp was obtained at the Smithsonian National Museum of Natural History with a Canon 60D DSLR, with a Canon MP-E 65 mm F/2.8 Macro photo lens and a Canon MT–24EX Macro Twin Lite Flash (Tokyo, Japan) with custom-made diffusers to minimise hot spots. Images were saved as TIF files and focus stacked using Zerene Stacker v. 1.04. Image editing was done in Adobe Photoshop, and plate layout was performed in Adobe Illustrator.

Previously collected expression data from various populations, organs, and life stages were used for annotation. For adult female venom glands and carcass (whole body tissue minus the venom gland) samples included the following: wasps collected from (a) wild low bush at Wolf Lake, MI (44.014027, −85.842384) in March 2023, (b) wild highbush at Otis Lake, MI (42.607586, −85.418655) in July 2021, and (c) two cultivated blueberries varieties, Jersey (42.4386, −86.217674) and Coville (43.392845, −84.24497) collected from farms in western and central MI, respectively, in July 2021. To obtain these samples, prior to adult emergence, galls were collected and immediately shipped overnight to the University of New Mexico. A subset of galls were individually placed in polystyrene *Drosophila* vials (Fisherbrand) and adult emergence was monitored daily. For larval salivary glands and malpighian tubules, galls were collected from a commercial farm in West Olive, Michigan (42.9715, −86.0765) in August 2021, which were immediately shipped overnight to the University of New Mexico where the larvae were immediately dissected out of the galls. See below for extraction and sequencing details.

### DNA library preparation and sequencing

The whole body of a single female wasp was homogenized into a fine powder while kept frozen using a SPEX SamplePrep 2010 Geno/Grinder (Cole Parmer, Metuchen, New Jersey, USA) and underwent high molecular weight (HMW) DNA extraction using the fresh or frozen tissue protocol of the Qiagen MagAttract HMW DNA Kit (Qiagen, Hilden, Germany). Following isolation, the HMW DNA (175ng) was sheared to a mean fragment length of 18 Kb with a Megaruptor 3 (Diagenode, Dennville, New Jersey, USA) and prepared into a PacBio SMRTBell library using the SMRTBell Express Template Prep Kit. 3.0 (Pacific Biosciences, Menlo Park, California, USA) using a barcoded adapter. After DNA isolation, shearing, and library preparation, the sample was purified using solid-phase reversible immobilization beads (SPRI beads) (DeAngelis et al. 1995) and quantified using fluorometry and spectrophotometric absorbance ratios (DeNovix Inc., Wilmington, Delaware, USA). Fragment length distributions after each step were determined by Femto Pulse or Fragment Analyzer (Agilent Technologies, Santa Clara, California, USA). The resulting library was pooled with other samples and sequenced on a PacBio Revio system using a 30-hour movie length on 1/4th of a Revio SMRTCell. Raw subreads were converted to HiFi data using the PacBio SMRTLink software v.26.1.0.284828.

Concurrent to HiFi sequencing, a pool of three females were used to prepare an enriched chromosome conformation capture (HiC) library. Tissues were homogenized in 1× phosphate buffered saline and nuclei were crosslinked in a 2% formaldehyde solution. Following crosslinking, the sample was lysed and digested using the restriction enzymes DdeI and DpnII. To enrich the sample for proximity ligated fragments, a biotin-labeled dATP fill-in step was performed prior to proximity ligation so fragments could be captured downstream. Following proximity ligation, a crosslink reversal step was performed followed by two DNA purification steps using SPRI beads, the removal of biotin from unligated ends, and another DNA purification step. The sample was size-selected using SPRI beads, biotinylated ligation products were captured, and the sample was prepared into a short-read sequencing library using the NEBNext Ultra II DNA Library Prep Kit (New England Biolabs, Ipswich, Massachusetts, USA). The final libraries were sequenced on a partial flow cell using the AVITI 2×150 Sequencing Kit Cloudbreak FS High Output kit on the Element AVITI System (Element Biosciences, San Diego, CA). Following sequencing, raw reads were basecalled using bases2fastq v.2.3.0.2116803307.

### Genome assembly, assessment, and contaminant removal

The HiFi reads were screened and filtered for adapter-contaminated sequence artifacts using FCS-Adaptor and HiFiAdapterFilt (Sim et al. 2022) (https://github.com/ncbi/fcs). The resulting HiFi reads were used to assemble contigs using HiFiASM (v.0.24.0-r702) (Cheng et al. 2021; Cheng et al. 2022) with implementation of the *–telo-m* flag with the canonical insect telomere sequence motif “CCTAA” (Vitkova et al. 2005). The contig assemblies were subsequently purged of duplicate contigs using PurgeDups (Guan et al. 2020), and the duplicate purged contig assembly served as the reference to map HiC reads using BWAmem 2 (v.2.2.1) (Vasimuddin et al, 2019). The resulting mapped reads were filtered for artifact PCR duplicates using Picard (v.3.2.0) (Picard2019toolkit, 2019 https://github.com/broadinstitute/picard). A contact map was generated from the de-duplicated mapped reads using the YaHS pipeline (Zhou et al. 2023). Visualization of the contact map and minor manual editing was achieved using Juicebox (v.2.15) (Durand et al. 2016). Minimap2 (v.2.22-r1101) was used to map HiFi reads back to the contig assembly and calculate coverage of each contig, the –auto function and –genome mode of BUSCO (v.6.0.0, Tegenfeldt et al. 2025) was used to select the appropriate taxon database and estimate genome completeness, with hymenoptera_odb12 (2025-07-01) as the selected lineage. BLAST+ and Diamond were used to perform nucleotide alignments to the NCBI nucleotide database (accessed November 2025) and UniProt protein database (accessed November 2025), respectively (Buchfink et al. 2021; Camacho et al. 2009; Li, 2018, Tegenfeldt et al. 2025). The resulting outputs of minimap2, BUSCO, BLAST+, and Diamond were summarized and visualized using Blobtools2 and blobblurb (Challis et al. 2020); (https://github.com/sheinasim/blobblurb). Additional taxonomic assignments of contigs and contig fragments was performed using the FCS-GX (Astashyn et al. 2024). The taxonomic assignments based on a combination nucleotide alignment, protein alignment, and FCS-GX were used to identify contigs assigned to *Wolbachia* and remove non-Arthropod contigs from the assembly. The primary and alternate assemblies were submitted to the National Center for Biotechnology Information (NCBI).

### RNA library preparation and sequencing

We sequenced the transcriptome of three tissue types of *H. nubilipennis*: venom gland dissected from adult females, the whole body of adult females after the venom gland was removed, as well as the Malphigian tubules and salivary glands dissected from late-instar larvae removed from galls. All dissections were performed under a dissecting microscope in individual drops of 30 µL ice-cold Gibco 1× phosphate-buffered saline (PBS) (ThermoFisher). The venom glands were dissected from adult female wasps using microforceps, and pools of 20 glands were collected for RNA extraction. From the same individuals, pools of five carcasses (all tissues excluding the venom glands) were also collected. Similarly, pools of 15 paired salivary glands and Malpighian tubules were dissected from actively feeding larvae. All samples were immediately placed into a lysis buffer from the Spectrum™ Plant Total RNA Kit (Sigma) and stored at −80 °C until extraction. RNA was extracted following the manufacturer’s protocol. THe quantity and quality of each RNA extraction was individually assessed prior to pooling using a NanoDrop 2000c (Thermo Scientific) and an Agilent 2100 Bioanalyzer. Library preparation and RNA sequencing were performed by Novogene. Raw sequencing reads were filtered and trimmed using Trimmomatic v0.36 (Bolger et al. 2014). Samples used for genome annotation included *H. nubilipennis* collected from four host sources: native lowbush (5 replicates of venom glands and carcasses); native highbush (4 replicates venom glands and carcasses); cultivated blueberry variety ‘Jersey’ (4 replicates of venom glands and carcasses, 6 replicates of larval salivary glands and Malpighian tubules, and 4 replicates of whole larvae), and cultivated blueberry variety ‘Coville’ (4 replicates of venom glands and carcasses). Expression values were obtained from Salmon quantification against the one-isoform-per-gene EGAPx transcript set (Patro et al. 2017).

### Genome Annotation

We performed genome annotation of *H. nubilipennis* using the Eukaryotic Genome Annotation Pipeline - External (EGAPx) (https://github.com/ncbi/egapx) v0.4.0. RNA data was aligned to the reference using STAR v0.15. Miniprot v2.7.11 was used to align Hymenoptera protein sequences to the reference. Gnomon (https://www.ncbi.nlm.nih.gov/refseq/annotation_euk/gnomon/) was used for gene prediction using protein and RNA-seq alignments and *ab-initio* predictions based on HMM. Lastly EGAPx adds functional annotations to the final structural annotation set based on orthology and model type and quality. Because the EGAPx annotation produced an unusually high number of predicted genes, we conducted a second annotation using BRAKER3 v3.0.8 (Gabriel et al. 2024) to independently evaluate gene predictions. The genome was first soft-masked with RepeatModeler and then annotated with BRAKER3 using the OrthoDB v12 Arthropoda protein set and one replicate from each RNA-Seq library (n = 32). Genomic elements were derived from the EGAPx annotation using bedtools v2.31.1 (Quinlan and Hall, 2010). Pfam domains were assigned using HMMER searches against the Pfam database (Eddy, 2011; Mistry et al. 2021).

### Methylome

During HiFi sequencing on the PacBio Revio, Jasmine v2.3.0 was used on-instrument to call 5-methylcytosine (5mC) modifications at CpG sites (Pacific Biosciences 2026a; Ni et al. 2023). Reads were aligned to the assembled chromosomes with pbmm2 v26.1.99 (align --preset CCS --sort) with MM/ML base modification tags preserved (Pacific Biosciences 2026b). CpG site methylation frequencies were estimated with pb-CpG-tools v3.0.0 (aligned_bam_to_cpg_scores, combined-strand, model mode) (Pacific Biosciences 2025; Ni et al. 2023). Sites with coverage <5 were excluded from analysis. A CpG site was considered methylated if its model-estimated methylation frequency exceeded 80%. For genomic element type comparisons, we calculated the fraction of callable CpG sites exceeding that threshold. Adapting the gene-body methylation criterion used for *Nasonia vitripennis* in Wang et al. (2013), a gene was considered methylated if it contained at least three callable exonic CpGs, of which at least 10% were methylated. Pfam domain enrichment among methylated genes was tested using a one-sided Fisher exact test against the eligible protein-coding background rate, with a Benjamini-Hochberg FDR correction.

### Mitochondrial genome

We identified all potential mitochondrial genome contigs using the MitoHiFi v3.2.3 pipeline (Uliano-Silva et al. 2023). MitoHiFi implemented a BLAST search for contigs that have a high similarity to the mitochondrial genome of *Sycobia* sp2 (NCBI RefSeq accession MT947600.2) (Wang et al. 2020) and selected the contig with the greatest similarity. Mitochondrial genes were then structurally annotated using intervals from the same mitochondrial genome used in the BLAST search through the MitFi annotation program in MITOS2 (Bernt et al. 2013). A representative mitochondrial genome was submitted to NCBI under the accession PZ709624. The cytochrome oxidase subunit I (COI) sequence fragment of our specimen were extracted and compared with the three *Hemadas* COI sequences available on GenBank: *H. nubilipennis* from Georgia (MW983184) and two *Hemadas* sp. in New York (MW983380, MW982580). All three GenBank sequences originate from specimens in the Smithsonian National Museum of Natural History collection (voucher numbers USNMENT01566055–57, Santos et al. 2023). We conducted the intraspecific divergence analysis using uncorrected *p*-distance using MEGA 12 (Kumar et al. 2024).

### *Wolbachia* endosymbiont

*Wolbachia* are maternally transmitted alpha proteobacteria (Rickettsiales) in many filarial worms and arthropods, and can induce a range of reproductive manipulation of their hosts ranging from cytoplasmic incompatibility, feminization of males, male killing, and induction of parthenogenesis in females (Vancaester & Blaxter, 2023; Fricke & Lindsey, 2024). The *Wolbachia* sequence was confirmed using BLAST (*wHem* henceforth), and annotated using Prokka v1.15.6 (Seemann 2014). A BLAST database was created for *wHem*, and we performed BLAST searches for parthenogenesis-inducing effector proteins PifA and PifB (Fricke & Lindsey, 2024). Finally, we assessed *wHem* using BUSCO lineage rickettsiales_odb12 (2025-05-14).

### Repeat landscape

We quantified the total repetitive DNA content of 16 chalcidoid assemblies across 11 families (**Table 2**) using 100% complete matrix of BUSCO output (2802 loci), the resulting gene trees were generated using BUSCO_phylogenomics pipeline (https://github.com/jamiemcg/BUSCO_phylogenomics), and the species tree (**Figure 3A**) was generated using ASTER v1.25 (Zhang et al 2025). Repeat annotation summaries generated using EarlGrey v7.1.0 (Baril et al. 2024) and visualized using custom R scripts (Zhang et al 2026, https://github.com/merondun/hymenopteran_alignments), assigning each genomic position to a single repeat subclass using a fixed hierarchical order (LTR, LINE, Penelope, SINE, DNA, Rolling Circle, Other, Unclassified, Non-Repeat), with overlapping intervals resolved by priority-based merging in this order using R, tidyverse, and GenomicRanges v1.56.2 (Lawrence et al. 2013). From these repeat annotations we extracted three summary metrics per transposable element (TE) subclass: (i) total cumulative genomic coverage (Mb), (ii) the number of distinct TE families, and (iii) the proportion of the genome covered by each subclass (**Figure 3B**). To assess the relationship between repeat abundance and genome size while accounting for shared evolutionary history, we performed phylogenetic generalized least squares (PGLS) regression using caper (v1.0.4) (Orme et al. 2025) in R. Both genome size and total repeat genomic coverage (Mb) were log-transformed and Pagel’s λ was estimated by maximum likelihood to identify phylogenetic signal. As a non-parametric complement, we computed Spearman’s rank correlation between genome size and repeat genomic coverage (Mb) (**Figure 3C**).

**Table 1.** Assembly statistics of the *Hemadas nubilipennis* genome contig and scaffold assemblies.

|  | Contig | Scaffold |
| --- | --- | --- |
| **Total Fragments** | 34 | 9 |
| **N50 (MB)** | 218 | 218 |
| **L50** | 7 | 3 |
| **N90 (MB)** | 189 | 189 |
| **L90** | 19 | 5 |
| **Total (MB)** | 1077.946 | 1077.948 |

**Table 2.**
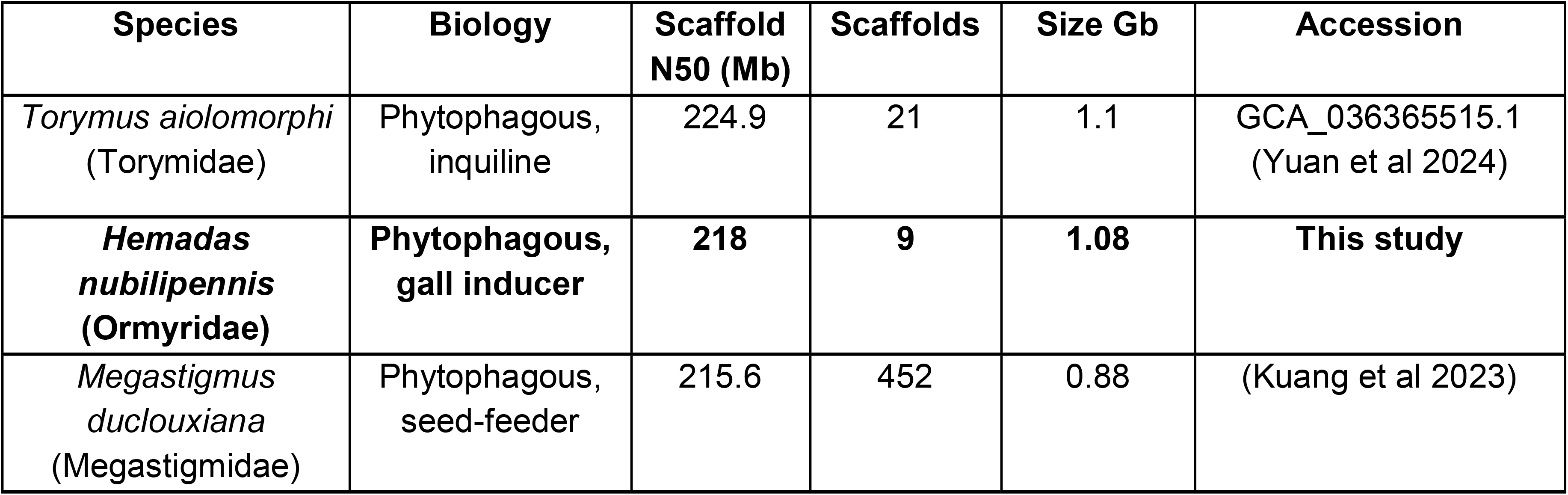
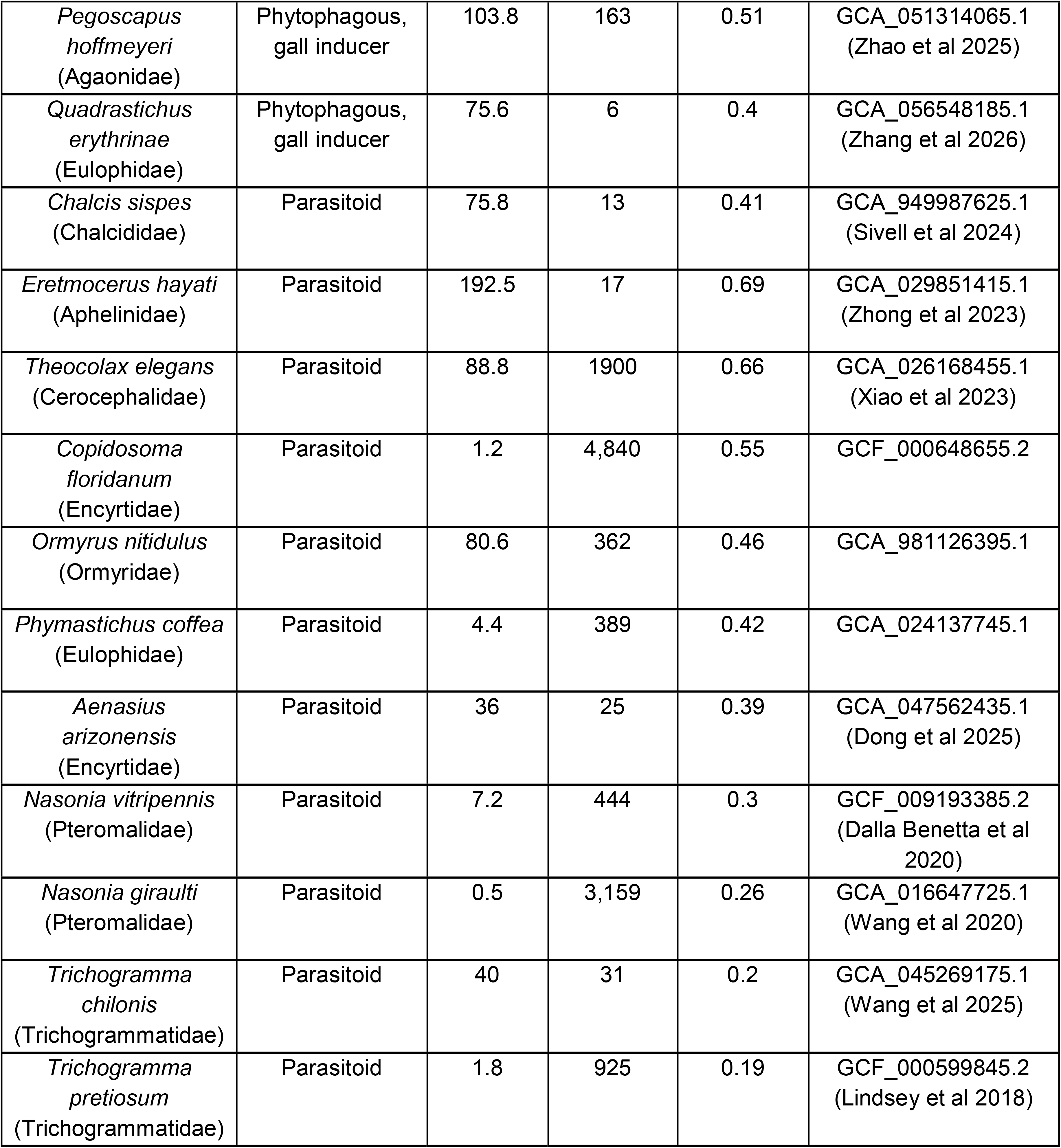
Assembly statistics and accession information for the chalcidoid genomes referenced for repeat evolution and/or methylation, sorted by biology then genome size.

## Results and Discussion

### Genome assembly and annotation

PacBio HiFi sequencing of *Hemadas nubilipennis* generated 1,705,747 reads (28.4 Gb) from one Revio cell, with 0.00146% discarded after filtering. Genome coverage was estimated at 25.6× by GenomeScope2. The initial HiFiASM assembly comprised 34 contigs (N50 = 218 Mb; L50 = 7; genome size = 1077.946 Mb; Table 1). Scaffolding with HiC improved contiguity, and BlobToolKit identified and removed one non-Arthropod scaffold corresponding to *Wolbachia pipientis*. The final assembly contains nine scaffolds totaling 1077.948 Mb (N50 = 218 Mb; L50 = 3; N90 = 189 Mb; L90 = 5; Figure 2A), with five chromosomes resolved by HiC (Figure 2B) and high homozygosity (Figure 2C). BUSCO completeness is C:94.6% [S:93.0%, D:1.6%], F:1.6%, M:3.8% (n = 5,920; Figure 2A). Quality metrics exceed Earth BioGenome Project standards (Lawniczak et al. 2022), including contig N50 > 1 Mb and >90% single-copy BUSCOs. At 1.077 Gb, this is the second largest chalcidoid genome after *Torymus aiolomorphi* (Yuan et al. 2024).

**Figure 2.**
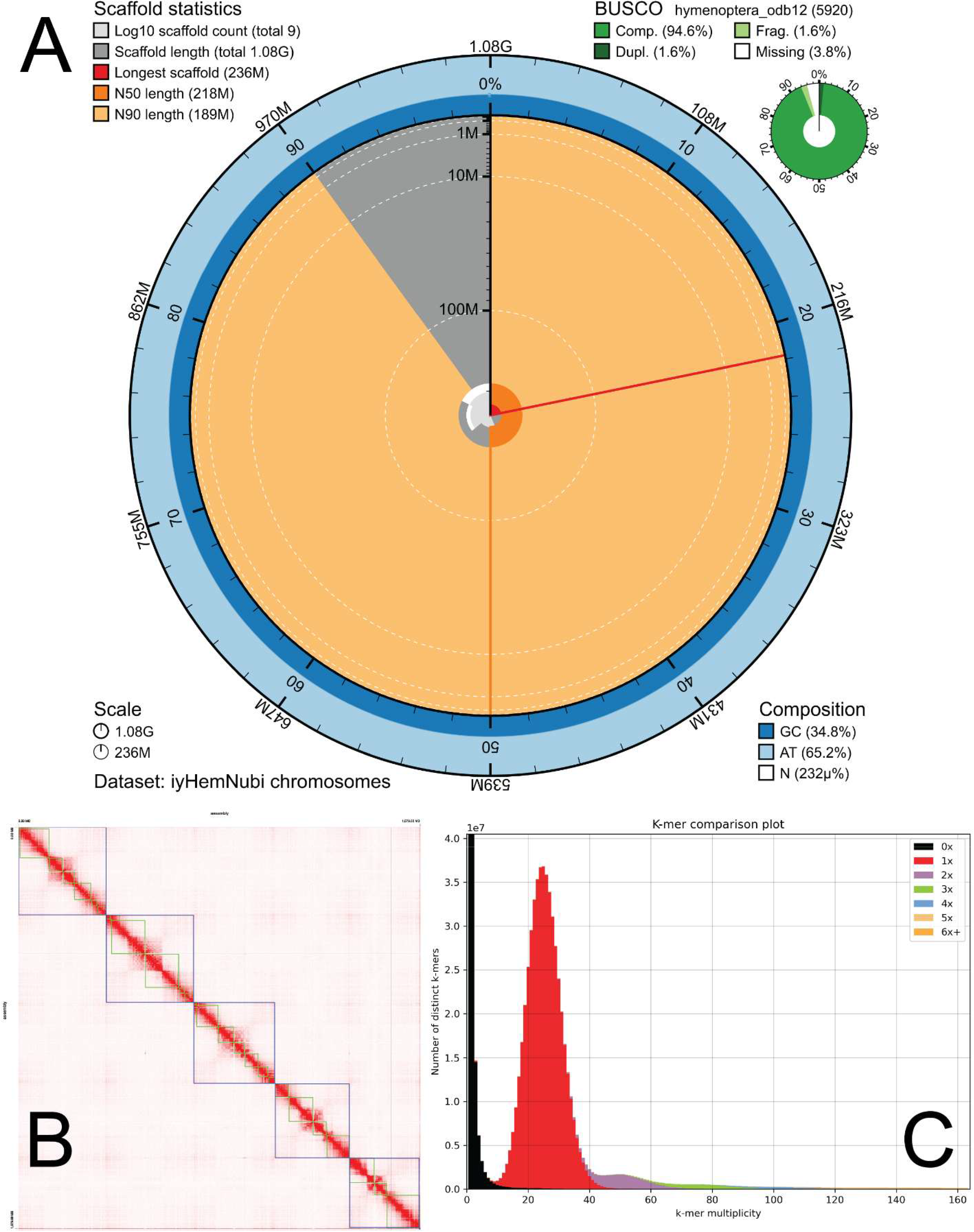
Statistics of the *Hemadas nubilipennis* assembly. **A)** Snail plot visualization of the scaffold assembly with BUSCO assessment using Hymenoptera conserved orthologs odb12. **B)** Hi-C contact map indicates that contigs can be grouped in 5 major scaffolds. **C)** K-mer spectra plot showing the representation graph of k-mers in the assembly relative to the raw HiFi data.

**Figure 3.**
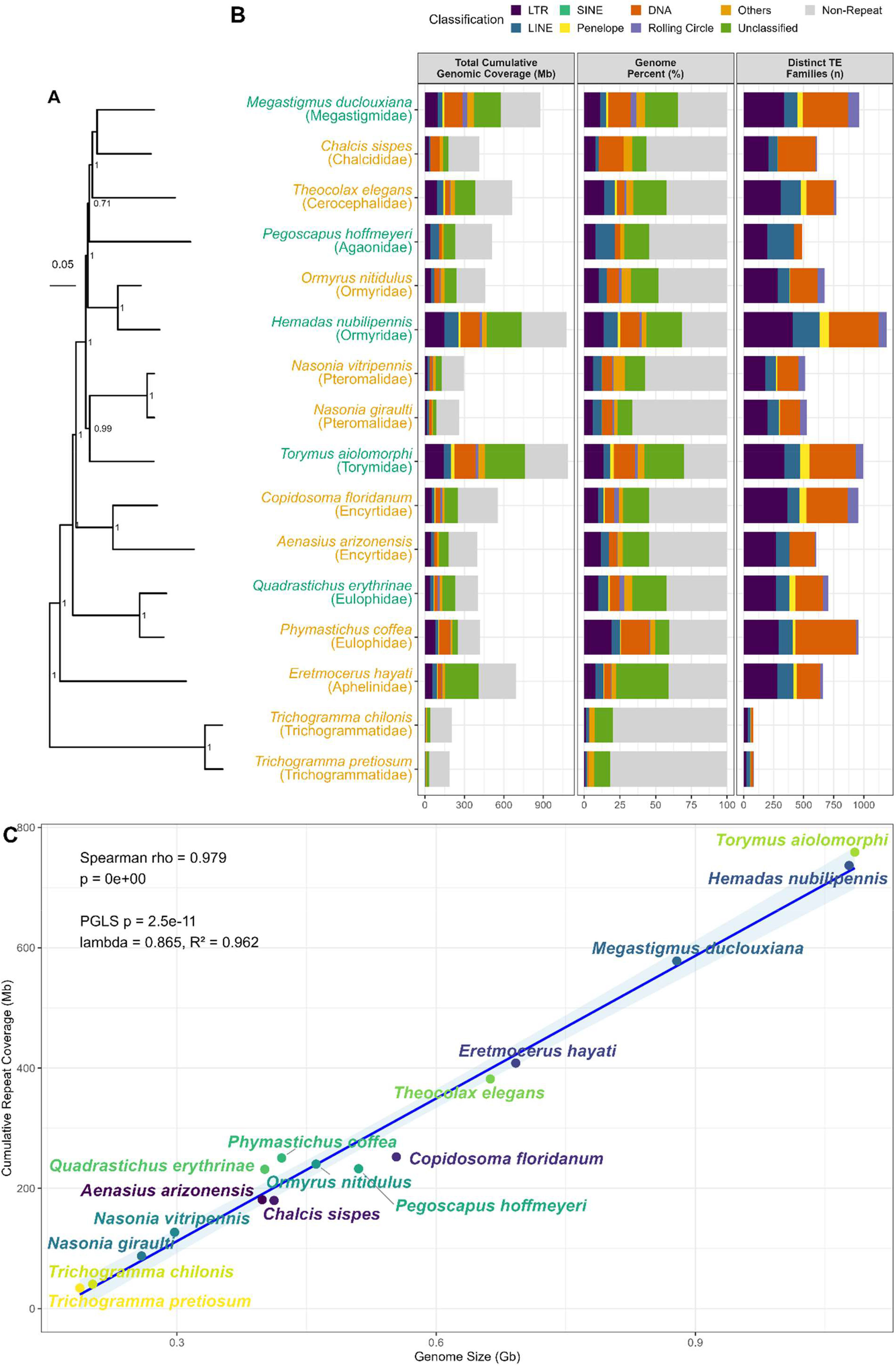
Repeat landscape across chalcidoid genomes. **A)** Phylogenetic relationship of the chalcidoid assemblies, with local posterior probability value for nodal support. Green tip label indicates phytophagous lineages, yellow indicates parasitoid. **B)** Total repetitive DNA content using EarlGrey and summarized by repeat subclass. Facets show total cumulative genomic coverage (Mb), percent of the genome occupied by each repeat subclass, and number of distinct transposable element (TE) families (“Non-Repeat”, “Unclassified”, and “Other” categories were excluded from TE family count visualizations). **C)** Association between genome size (Gb) and cumulative repeat coverage (Mb) with PGLS regression and Spearman correlation results annotated.

Repetitive elements comprise 68.7% (740.3 Mb) of the genome: class I TEs (LTRs, LINEs, SINEs, Penelope-like) account for 24.9%, class II TEs (DNA transposons, Rolling Circle helitrons) for 15.3%, unclassified TEs for 25.0%, and other repeats (simple repeats, microsatellites, RNAs) for 3.5%.

EGAPx annotated 31,818 genes (25,023 protein-coding, 6,795 lncRNA), whereas BRAKER3 predicted 18,405 gene models (25,698 isoforms) with similar BUSCO scores (C:95.2% [S:93.7%, D:1.6%], F:0.7%, M:4.0%). However, RNA-Seq read mapping (109 libraries) averaged 74.8% for EGAPx versus 46.1% for BRAKER3, indicating that EGAPx captured many RNA-supported genes missed by BRAKER3.

### Gene body methylation

*Dnmt3*, encoding a de novo DNA methyltransferase, is retained in *H. nubilipennis*, consistent with its presence in congeners *Ormyrus nitidulus* and *O. pomaceus* (Kucharski et al. 2023). Methylation is strongly enriched in exons: 1.94% of exonic CpG sites were methylated (>80% 5mC), versus 1.02% in promoters, 0.13% in introns, and 0.06% in intergenic regions (total CpG sites = 39,282,200; 92,623 methylated, 0.24%). Only 3,182 of 24,979 protein-coding genes (12.7%) with ≥3 callable exonic CpGs had ≥10% methylation. Among methylated genes, methylation was low in exon 1 (5.3%), peaked in exons 2–4 (39–51%), and tapered thereafter. These genes are enriched for Pfam domains associated with core cellular functions (WD40/β-propeller, TPR proteins, translation, RNA processing, ubiquitin-proteasome components), mirroring patterns in *Nasonia vitripennis* and other chalcidoids (Wang et al. 2013, 2015; Bewick et al. 2017; Lindsey et al. 2018).

Methylated genes show higher median expression than unmethylated genes (19.11 vs. 0.68 TPM) and broader tissue distribution (83.6% vs. 19.9% expressed across all five sample types). Methylated CpG density correlates positively with gene-rich, BUSCO-rich chromosome arms (Spearman ρ = 0.60 over 100-kb windows) and approaches zero in gene-poor centromeric regions (Figure 4). These findings indicate that *H. nubilipennis* possesses a conserved chalcidoid gene-body methylation profile.

**Figure 4.**
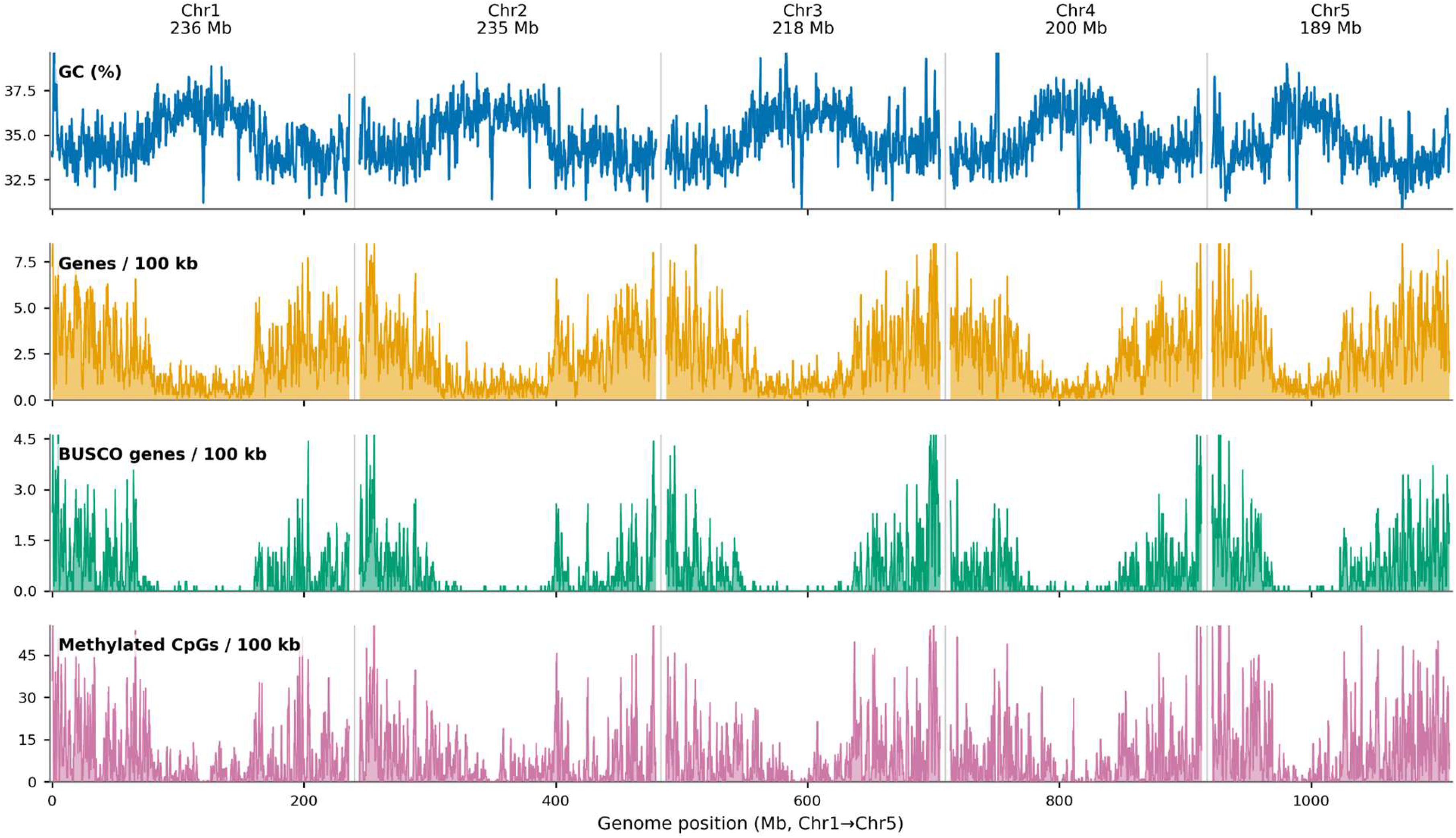
Chromosome structure of *Hemadas nubilipennis*. Tracks computed in 100-kb windows along each of five chromosome-scale scaffolds, smoothed with a 700-kb rolling mean. From top to bottom, per 100 kb: GC content (%); EGAPx-predicted protein-coding gene density; conserved single-copy gene density (genome-mode BUSCO hymenoptera_odb12 “complete” genes); density of methylated CpG sites (>80% 5mC).

### Mitochondrial genome

The mitochondrial genome is 17,270 bp and contains 13 protein-coding genes, 21 tRNAs (missing *trnH*), and both rRNAs. Gene order (cox1–cox2–atp8–atp6–cox3–nad3–nad2–rrnS–rrnL–nad1–cytb–nad6– nad4L–nad4–nad5) differs from the type-5/6 arrangements common in Chalcidoidea (Zhu et al. 2023): the cox/atp/nad3 block is inverted (type-5/6: nad3–cox3–atp6–atp8–cox2–cox1; *H. nubilipennis*: cox1–cox2– atp8–atp6–cox3–nad3), the nad5–nad4–nad4L block is reversed, and nad2 is relocated from its terminal position in type-5/6 to an early position. This extensive rearrangement resembles that of the distantly related gall-inducer *Quadrastichus erythrinae* (Zhang et al. 2026).

COI sequences from our Michigan specimen (lowbush blueberry) show 4.3% divergence from a Georgia specimen (highbush) and 5.4% from New York specimens (unknown hosts); Georgia and New York differ by 2.8%. These genetic distances, combined with host plant differences, subtle morphological variation (L. F. Nastasi, pers. comm.), gall morphology variation, and sex ratio differences, suggest that *Hemadas* may represent a cryptic species complex or multiple host races. The holotype (Ashmead 1887) from Toronto, Canada, suggests our Michigan specimen may represent the true *H. nubilipennis*, but a comprehensive taxonomic revision is needed.

### *Wolbachia* endosymbiont

The *Wolbachia* genome wHem (1,376,038 bp) has a BUSCO score of 98.3% single-copy complete, 1.2% fragmented, 0% duplicated, and 0.6% missing (rickettsiales_odb12, n=345), with 1,457 CDS, 3 rRNA, 34 tRNA, and 1 tmRNA. It belongs to Supergroup A, common in terrestrial insects including Hymenoptera (Vancaester and Blaxter 2023). wHem encodes PifA and PifB effector proteins, which may induce parthenogenesis (Fricke and Lindsey 2024). Wolbachia-induced cytoplasmic incompatibility could also promote reproductive isolation between populations, potentially facilitating cryptic speciation among lowbush and highbush hosts, analogous to its role in the cryptic species complex of *Cotesia* parasitoid wasps (Valerio et al. 2024).

### Repeat evolution across Chalcidoidea

The BUSCO-based phylogeny places *H. nubilipennis* as sister to *Ormyrus nitidulus* (Figure 3A), supporting the revised Ormyridae classification (van Noort et al. 2024; Hanson et al. 2025). Repeat composition varies widely across 16 chalcidoid genomes (Figure 3B). Total repeat content is strongly correlated with genome size (Spearman’s ρ = 0.979, p < 0.001), and PGLS regression confirms this relationship after accounting for phylogeny (λ = 0.865, R² = 0.962, p = 2.5e−11), explaining ∼96% of size variance (Figure 3C).

Phytophagous chalcidoid genome sizes range from 1.1 Gb (*T. aiolomorphi*; Yuan et al. 2024) to 0.4 Gb (*Q. erythrinae*; Zhang et al. 2026). Phytophagous species have larger genomes than parasitoid relatives in Ormyridae (*H. nubilipennis* vs. *O. nitidulus*), but not in Eulophidae (*Q. erythrinae* at 399.2 Mb vs. *P. coffea* at 421.2 Mb; Figure 3B). DNA transposons and unclassified TEs dominate the TE landscape in *H. nubilipennis* and many phytophagous lineages, though some parasitoids also show this pattern (e.g., *Chalcis sispes* and *P. coffea* have high DNA transposon content; *Eretmocerus hayati* has abundant unclassified TEs; Figure 3B).

The 1.08-Gb genome of *H. nubilipennis* is driven predominantly by TE expansion (68.3%), consistent with other chalcidoid studies (Ye et al. 2022; Zhong et al. 2026). The strong TE-genome size relationship supports the hypothesis that phytophagy, especially gall induction, may promote genome expansion in Hymenoptera, though exceptions (Eulophidae) indicate lineage-specific effects. The elevated proportion of unclassified TEs in *H. nubilipennis* and other phytophagous lineages suggests a dynamic, potentially convergent expansion (Ye et al. 2022; Zhong et al. 2026). TE activity can generate regulatory novelty and facilitate rapid adaptation to selection (Serrato-Capuchina and Matute 2018), possibly enabling *H. nubilipennis* to shift among native lowbush, native highbush, and cultivated highbush hosts. The recent outbreak in Michigan cultivated highbush, despite the species’ native status and historically low densities, raises the possibility that TE-mediated genomic changes facilitated adaptation to anthropogenically modified environments. Population genomic comparisons across host associations, enabled by this reference genome, are needed to test this hypothesis.

### Implications for pest management

This genomic resource provides tools for addressing the BSGW outbreak. A rapid COI-based diagnostic would enable monitoring of genetic lineages that may differ in pest status, host preference, or phenology (Doellman et al. 2020). This is critical because the Michigan outbreak is restricted to cultivated highbush, whereas lowbush and native highbush populations remain at low densities. The wHem genome also offers potential for biological control via symbiont manipulation.

On the plant side, four QTLs associated with galling resistance have been identified in northern highbush blueberry, with candidate genes linked to defense, biotic stress, and phytohormone metabolism (Teresi et al. 2025). Resistant cultivars like ‘Draper’ mount an early defense that kills eggs within days, whereas susceptible genotypes like ‘Liberty’ show delayed responses permitting gall development. These four loci jointly explain 70.2% of the genetic contribution to resistance (Teresi et al. 2025). Integrating these plant-side findings with the wasp genomic resources creates a framework for understanding the molecular interaction between *H. nubilipennis* and its blueberry hosts.

## Conclusion

This comparative genomic analysis demonstrates that chalcidoid genome size is overwhelmingly governed by transposable element load, with *H. nubilipennis* carrying one of the largest repertoires known. The near-complete *Wolbachia* genome and conserved methylome provide immediate molecular targets for investigating parthenogenesis and host adaptation. Critically, COI divergence among populations suggests that what is currently treated as a single pest species likely comprises a cryptic complex, underscoring the need for population genomics to inform taxonomy and management. Integration of these genomic resources with ongoing blueberry resistance research will be essential for developing sustainable management strategies for BSGW.

## Data availability

The primary and alternate genome assemblies are hosted at the National Center for Biotechnology Information (NCBI) under BioProject Accessions PRJNA1484684 and PRJNA1484697 respectively. The sample used for the contig assembly is described under BioSample SAMN61252714 and registered under the Darwin Tree of Life ID iyHemNubi1. The RNAseq data used for annotation is under PRJNA1511513. The mitogenome is under the GenBank accession number PZ709624, and the *Wolbachia* wHem is under SAMN61279751. Raw read data was submitted under SRA accessions: SRR39418079–80.

## Acknowledgments

We thank Michael Gates (USDA-SEL) for access to the specimen and imaging system at the Smithsonian National Museum of Natural History, and Louis Nastasi (University of Nebraska-Lincoln) for discussions on the morphology of *Hemadas* species. YMZ is supported by Oak Ridge Institute for Science and Education (ORISE) fellowship, This research used resources provided by the USDA-ARS project number 2040-30400-003-000D, USDA-ARS SCINet project numbers 0201-88888-003-000D and 0201-88888-002-000D, USDA-NIFA accession 1031309, NSF Award Number 2305880, and the Smithsonian Institution High Performance Cluster (https://doi.org/10.25572/SIHPC). GRH was supported by NSF award number 2418250.

## Conflicts of interest

The authors declare that the research was conducted in the absence of any commercial or financial relationships that could be construed as a potential conflict of interest. All opinions expressed in this paper are the authors’ and do not necessarily reflect the policies and views of USDA. Mention of trade names or commercial products in this publication is solely for the purpose of providing specific information and does not imply recommendation or endorsement by the U.S. Government. USDA is an equal opportunity provider and employer. The authors declare no conflict of interest.

---

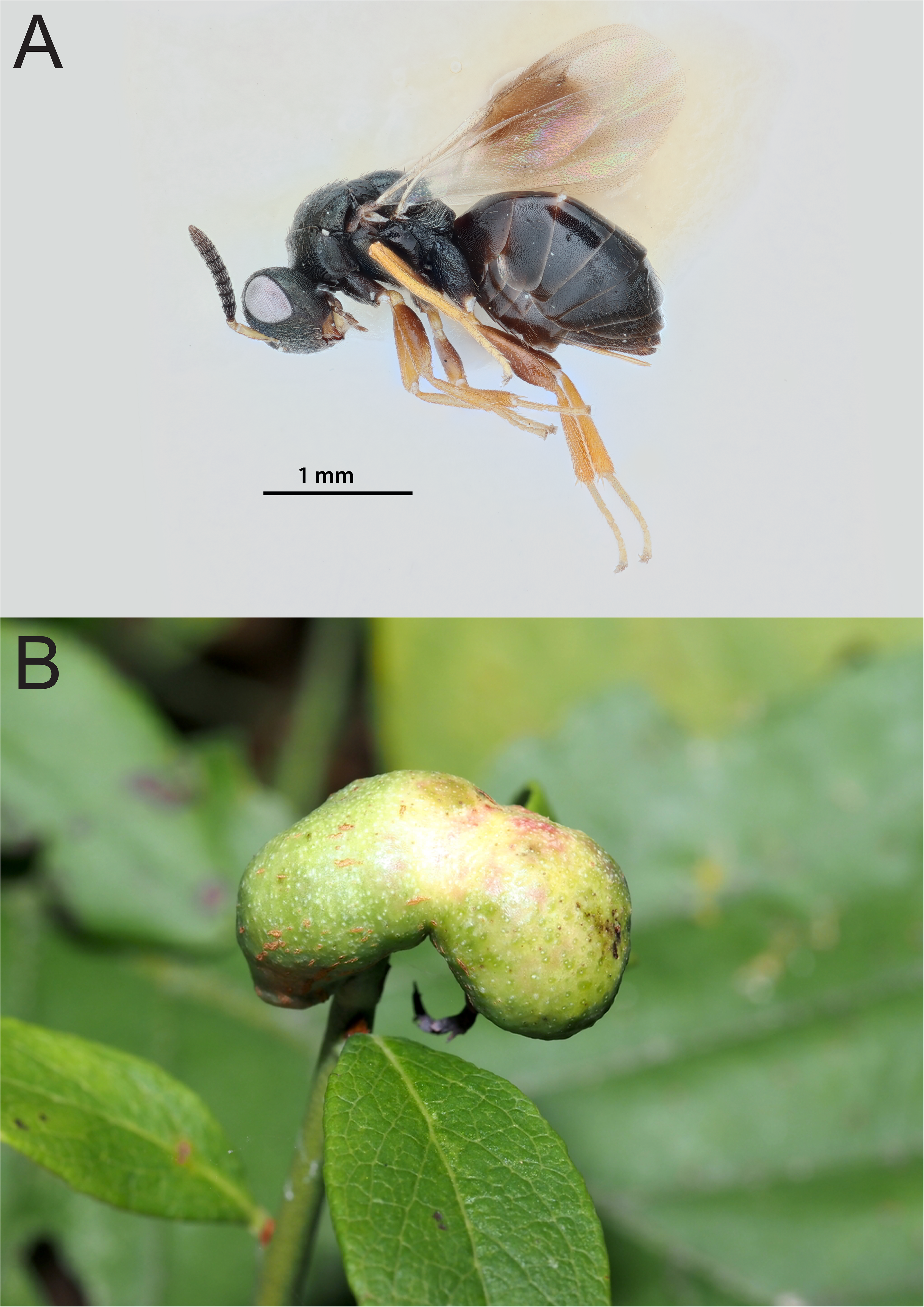

